# TranslucentID: Detecting Individuals with High Confidence in Saturated DNA SNP Mixtures

**DOI:** 10.1101/390146

**Authors:** Darrell O. Ricke, James Watkins, Philip Fremont-Smith, Tara Boettcher, Eric Schwoebel

## Abstract

High throughput sequencing (HTS) of complex DNA mixtures with single nucleotide polymorphisms (SNPs) panels can identify multiple individuals in forensic DNA mixture samples. SNP mixture analysis relies upon the exclusion of non-contributing individuals with the subset of SNP loci with no detected minor alleles in the mixture. Few, if any, individuals are anticipated to be detectable in saturated mixtures by this mixture analysis approach because of the increased probability of matching random individuals. Being able to identify a subset of the contributors in saturated HTS SNP mixtures is valuable for forensic investigations. A desaturated mixture can be created by treating a set of SNPs with the lowest minor allele ratios as having no minor alleles. Leveraging differences in DNA contributor concentrations in saturated mixtures, we introduce TranslucentID for the identification of a subset of individuals with high confidence who contributed DNA to saturated mixtures by desaturating the mixtures.

## Introduction

Panels of SNPs can be specifically designed to identify multiple individuals in complex DNA samples^1-4^. Isaacson et al.^2^ introduced SNP panel mixture analysis with drop-in and drop-out alleles present with confidence calculations for detecting individuals in mixtures. The confidence with which an individual can be identified in a DNA mixture using a given SNP panel is dependent upon the number of SNP loci in the panel and the average population ratio of minor to major SNP alleles^1,2^. Ricke and Schwartz^5^ refined the calculations for larger HTS SNP panels. The ability of SNP panels to differentiate contributors from non-contributors is impacted when the number of DNA contributors saturate the mixture (e.g., a mixture with detected minor alleles at a majority of the SNP loci).

Current DNA forensics relies upon the analysis of forensic samples by sizing short tandem repeats (STRs) with capillary electrophoresis. STR stutter alleles^6^ created by the polymerase chain reaction (PCR) loss and addition of STR repeat units limit the utility of analysis of mixtures by STR sizing. STR artifacts cannot be distinguished from alleles of low DNA contributor. Shifting from sizing STR alleles to HTS sequencing them^7-9^ will enable resolution of different STR alleles with identical lengths, but PCR allele stutter artifacts will remain. Because of the importance of analyzing complex DNA forensic mixtures, multiple solutions have tackled DNA mixtures of more than two mixtures: ArmedXpert^10^, DNAmixtures^11^, DNA VIEW Mixture Solution^12^, GeneMarker HID^13^, FST^14^, GenoProof Mixture^15^, GenoStat^16^, Lab Retriever^17^, LiRa^18^, LiRaHt, LRmix Studio^19^, MixSep^20^, STRmix^21^, and TrueAllele^22^. However, the current dogma is to avoid analysis of mixtures with more than two contributors as the results from multiple investigators on the same samples are discordant^23^.

DNA sources can vary by forensic samples. For touched samples, varying amounts of DNA are deposited on surfaces and objects by skin contact with individuals^24^. The amount of DNA varies by individual, object touched, how long the object was touched, DNA shedder level, and other factors. Hence, the amount of DNA contributions to forensic samples likely varies by individual for touched objects. We reason that the higher DNA contributors to saturated mixtures can be detected in saturated mixtures by treating a subset of the SNP loci with the lowest levels of detected minor alleles as likely major alleles for these higher DNA contributors. Both high and lower DNA contributors may be detectable as the saturation level is decreased by desaturation. We introduce TranslucentID, a new method to analyze saturated DNA mixtures to identify contributors to saturated mixtures with high confidence (patent pending)^25^.

## Methods

Multiple individuals can be identified in HTS sequenced SNP DNA mixtures with high confidence^2,3^. The confidence of such identifications can be quantified by the probability that a random man not excluded, or P(RMNE)^2,5^. Since saturated mixtures contain a lower number of major allele loci, it is less likely to identify individuals in saturated mixtures with low P(RMNE) using standard mixture analysis techniques. By increasing the number of loci called with no minor alleles in the mixture for comparison of the mixture with references, it is possible to identify individuals with low P(RMNE) in saturated mixtures. Individual reference profiles can then be matched to this desaturated mixture using standard mixture analysis techniques.

### Saturation level

HTS sequenced mixture and reference profiles often have a very small set of SNP loci with called allele counts below a minimum analytical threshold to rely upon the observed alleles. The MIT Lincoln IdPrism system uses a minimum of 50 called reads for all analyzed reference and mixture profiles. As a result, the number of compared SNPs between a mixture and each reference can vary to a small extent. Hence, the calculated saturation level for a mixture can vary slightly for each reference comparison; some comparison saturation levels will be higher than the panel saturation level. To avoid false positives, the IdPrism system uses a comparison saturation level upper limit of 70% (calculated by FastID).

### Implementation

The bottom k minor alleles by minor allele count or minor allele ratio (mAR) in the saturated mixture are treated as major alleles, such that the resulting desaturated mixture has N% major alleles. Equation 1 defines the value of k for a saturated mixture with M major allele positions sequenced on a panel of P total loci.

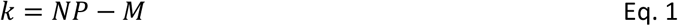

### Mixture Desaturation

A HTS mixture can be desaturated by treating the SNPs with the lowest mAR as mixture major alleles. As the number of mixture major alleles increases, the reference comparison results vary by contributed DNA concentration amount. For high DNA contributors, the number of dropped alleles will remain very low and the calculated P(RMNE) values will increase in significance. For intermediate DNA contributors, the P(RMNE) significance levels will peak at a value of k unique to that comparison as a function of increasing major alleles and the number of dropped alleles. For lower DNA contributors, the number of dropped alleles will increase linearly with increased k and P(RMNE) values will decrease with increased k. As k increases, the comparison saturation levels may drop below the upper limit enabling individual identification in the desaturated mixture.

### Matches to First Degree Relatives in Mixtures

First degree relatives share approximately 50% of their DNA. Parents and children share an allele at each autosomal SNP locus. For parents and children, the expected number of minor allele differences is defined by equation 1. This models the expected rate of observing a minor allele at k SNP loci.

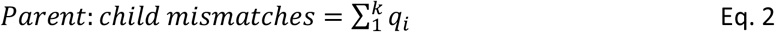

On average, siblings share two alleles at 25% of the autosomal loci plus one allele at 50% of the autosomal loci. For siblings, the expected number of minor allele differences is defined by equation 3. This models the expected rate of observing one or two minor alleles at the 25% of k loci with no shared alleles plus the expected rate of observing a minor allele at the 50% of k loci with one shared major allele.

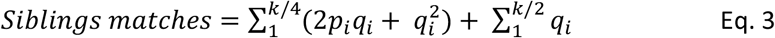

### Matches to Unrelated Individuals in Mixtures

An individual whose DNA does not contribute to a mixture and who is not related to any DNA contributes will have an expected number of mismatches defined by equation 4 for a mixture with k major allele loci.

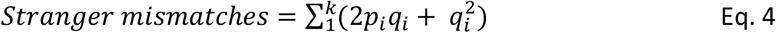

### Ion Torrent HTS Sequencing

The Ion Torrent HTS sequencing protocol described in Ricke et al.^26^ was followed.

### HTS SNP Data Analysis

The GrigoraSNPs^27^ program was used to call SNP alleles from multiplexed HTS FASTQ sequences. Mixture analysis was performed using FastID^3^ and Fast P(RMNE)^5^ as part of the MIT Lincoln Laboratory IdPrism HTS DNA Forensics system.

## Results

The mAR profiles for the eleven mixtures are illustrated in Figure 1. IdPrism uses a lower mAR analytical threshold of 0.001 to call minor alleles in mixtures. Below mAR of 0.001, SNP loci with random sequencing errors are observed in reference profiles^26^.

**Figure 1:**
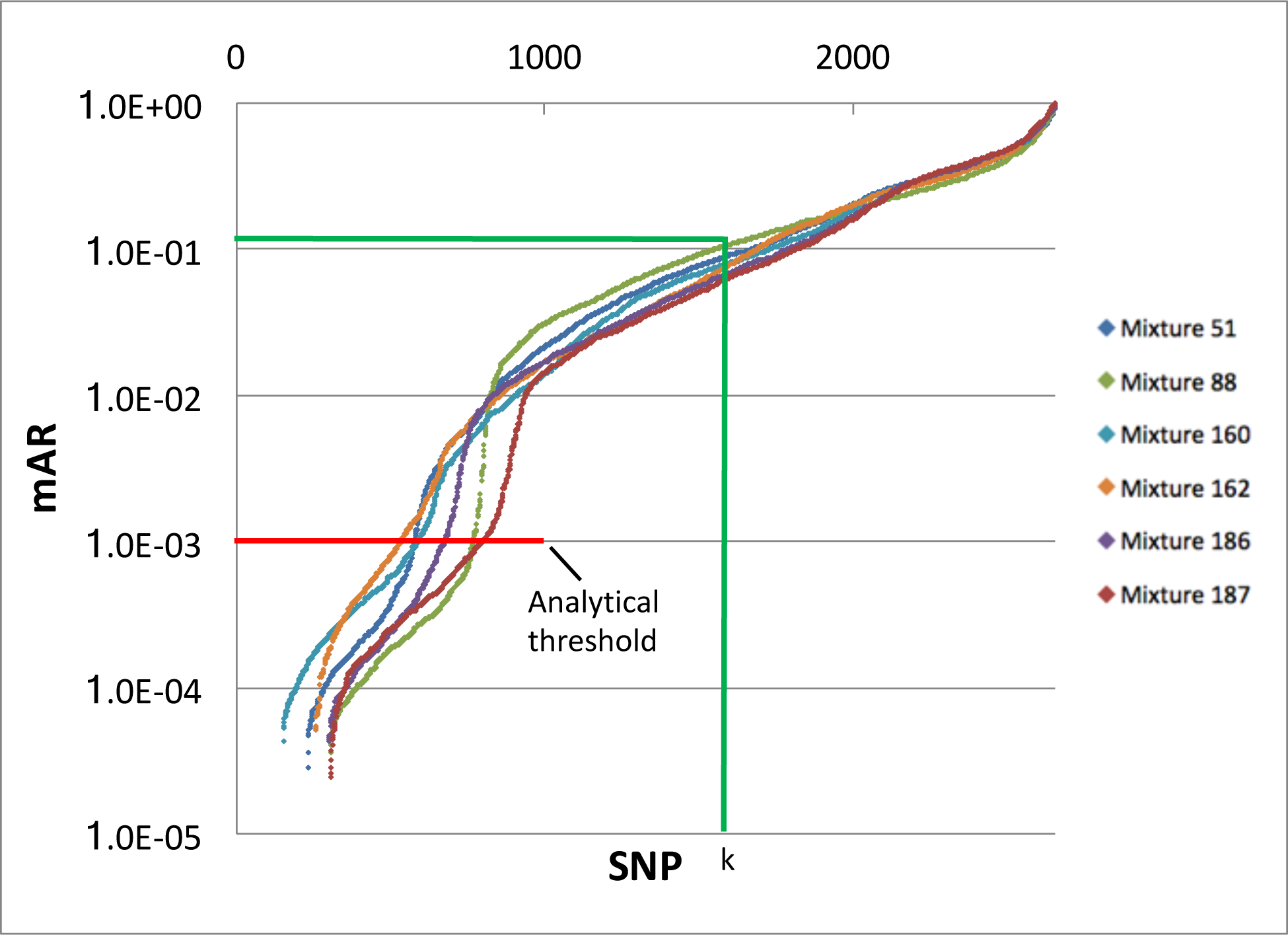
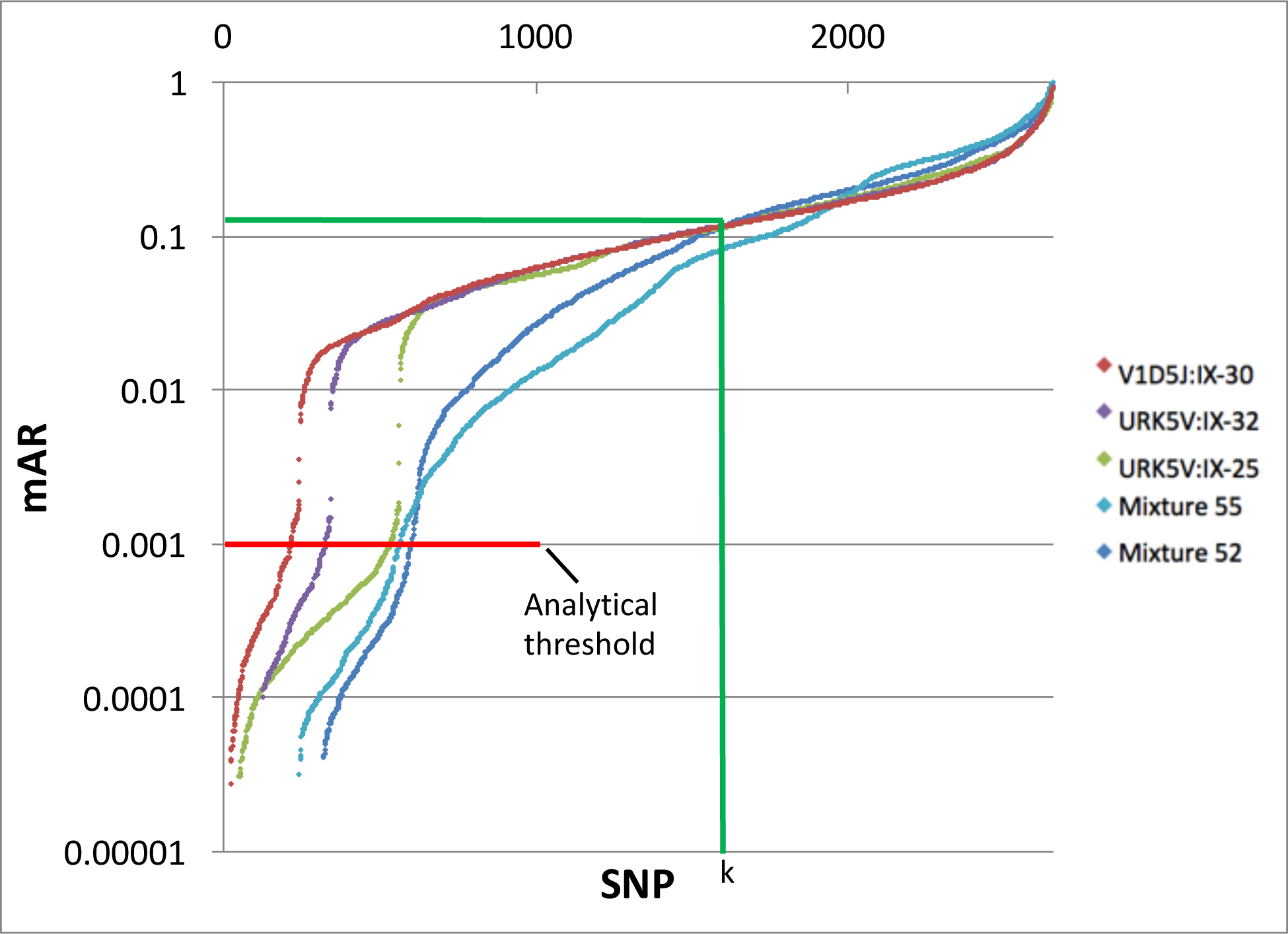
The eleven saturated mixtures used to demonstrate mixture desaturation. Loci are plotted in order of increasing mAR for each mixture.

Three equimolar mixtures of DNA with 10, 15, and 20 individuals were identified as saturated with minor alleles called at greater than 70% of the SNP loci. A set of 215 HTS mixture samples were examined to identify 8 saturated mixtures from touch samples. The number of SNP loci with called major alleles varies for each of these 11 mixtures (Figure 1). In the touch sample experiments, controlled sets of volunteers touched specific objects over the course of several days. Each object was touched one or more times on different days; some volunteers touched the same objects on multiple days. The order of touching objects and the set of volunteers touching the objects varied for each object. In these 11 saturated mixtures, mixture analysis with FastID^3^ identified only one individual in three mixtures (mix51, mix160, and mix186), and two individuals in (mix52, mix88, and mix187), see Table 1; in the remaining five mixtures, standard mixture analysis failed to detect any contributors. With the application of the new desaturation method, TranslucentID, an additional 41 individuals were identified in these mixtures with P(RMNE) values ranging from 2.70e-9 to as low as 8.39e-83 (mix55). The number of additional individuals identified in these mixtures range from one in mix88 and mix187 to as high as 9 for URK5V:IX-25.

**Table 1:**
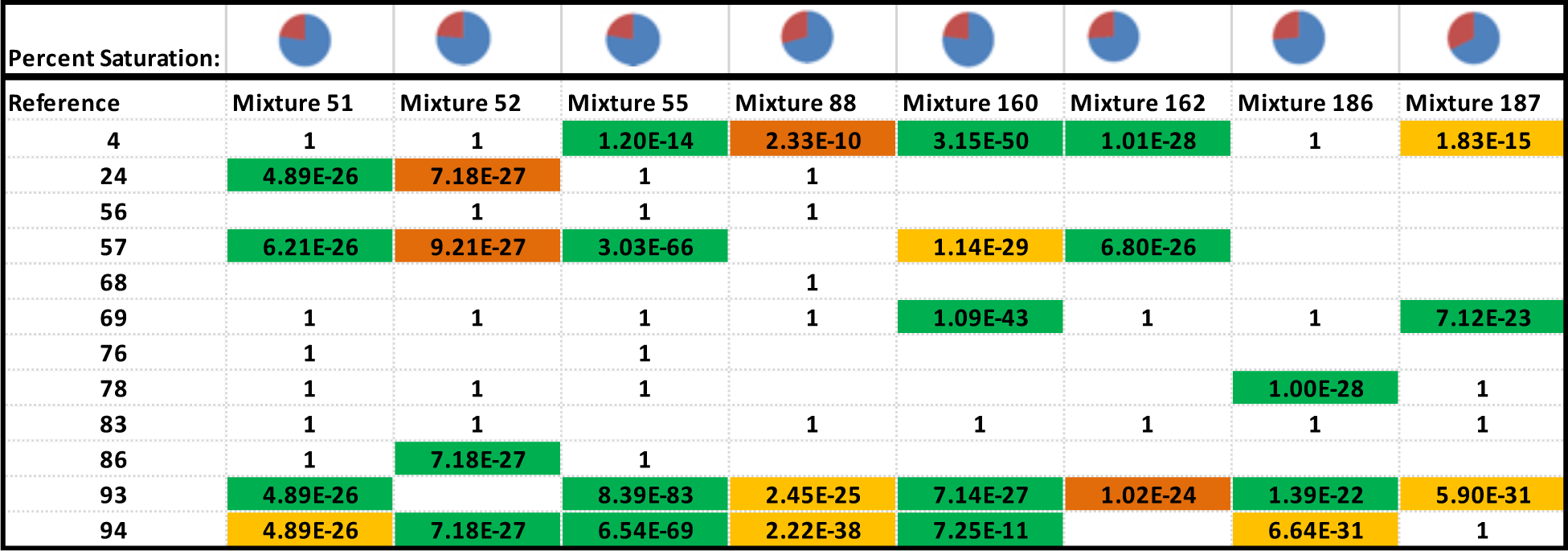
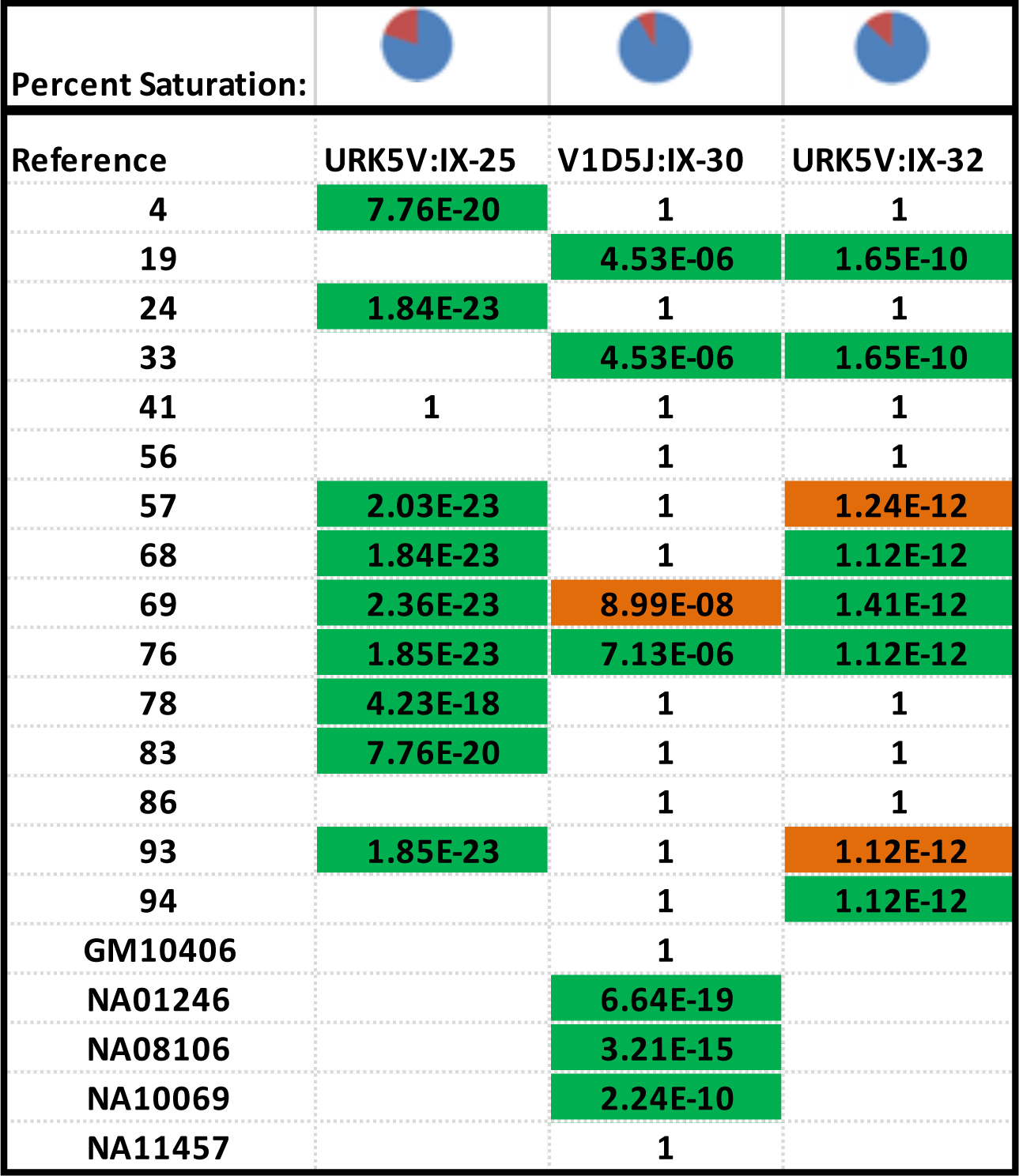
Performance of N=45% mixture desaturation. Mixtures are shown along the top of the table, and the pie charts represent the percent saturation of each mixture, where the blue sector represents the loci with one or more minor alleles detected and the red sector represents the loci with no called minor alleles. Green cells show new reference to mixture hits detected with TranslucentID desaturation, and the cells contain the probability that the random man is not excluded (PRMNE) of the match. The orange cells contain reference to mixture hits that were identified using both current mixture analysis and mixture desaturation.

| Percent Saturation: |  |  |  |  |  |  |  |  |
| --- | --- | --- | --- | --- | --- | --- | --- | --- |
| Reference | Mixture 51 | Mixture 52 | Mixture 55 | Mixture 88 | Mixture 160 | Mixture 162 | Mixture 186 | Mixture 187 |
| 4 | 1 | 1 | 1.20E-14 | 2.33E-10 | 3.15E-50 | 1.01E-28 | 1 | 1.83E-15 |
| 24 | 4.89E-26 | 7.18E-27 | 1 | 1 |  |  |  |  |
| 56 |  | 1 | 1 | 1 |  |  |  |  |
| 57 | 6.21E-26 | 9.21E-27 | 3.03E-66 |  | 1.14E-29 | 6.80E-26 |  |  |
| 68 |  |  |  | 1 |  |  |  |  |
| 69 | 1 | 1 | 1 | 1 | 1.09E-43 | 1 | 1 | 7.12E-23 |
| 76 | 1 |  | 1 |  |  |  |  |  |
| 78 | 1 | 1 | 1 |  |  |  | 1.00E-28 | 1 |
| 83 | 1 | 1 |  | 1 | 1 | 1 | 1 | 1 |
| 86 | 1 | 7.18E-27 | 1 |  |  |  |  |  |
| 93 | 4.89E-26 |  | 8.39E-83 | 2.45E-25 | 7.14E-27 | 1.02E-24 | 1.39E-22 | 5.90E-31 |
| 94 | 4.89E-26 | 7.18E-27 | 6.54E-69 | 2.22E-38 | 7.25E-11 |  | 6.64E-31 | 1 |

|  |  |  |  |
| --- | --- | --- | --- |
| **Percent Saturation:** |  |  |  |
| **Reference** | **URK5V:IX-25** | **V1D5J:IX-30** | **URK5V:IX-32** |
| 4 | 7.76E-20 | 1 | 1 |
| 19 |  | 4.53E-06 | 1.65E-10 |
| 24 | 1.84E-23 | 1 | 1 |
| 33 |  | 4.53E-06 | 1.65E-10 |
| 41 | 1 | 1 | 1 |
| 56 |  | 1 | 1 |
| 57 | 2.03E-23 | 1 | 1.24E-12 |
| 68 | 1.84E-23 | 1 | 1.12E-12 |
| 69 | 2.36E-23 | 8.99E-08 | 1.41E-12 |
| 76 | 1.85E-23 | 7.13E-06 | 1.12E-12 |
| 78 | 4.23E-18 | 1 | 1 |
| 83 | 7.76E-20 | 1 | 1 |
| 86 |  | 1 | 1 |
| 93 | 1.85E-23 | 1 | 1.12E-12 |
| 94 |  | 1 | 1.12E-12 |
| GM10406 |  | 1 |  |
| NA01246 |  | 6.64E-19 |  |
| NA08106 |  | 3.21E-15 |  |
| NA10069 |  | 2.24E-10 |  |
| NA11457 |  | 1 |  |

## Discussion

The number of contributors and the amount of contributed DNA can vary widely in saturated mixtures. The individuals detected by mixture analysis varies for mixtures by the amount of desaturation used, see Table 1. For detectable individuals, there tends to be a percentage of desaturation that has the lowest P(RMNE) value for each contributor. Apparent lower DNA contributors P(RMNE) values will be less significant as the desaturation amount increases due to increasing number of dropped alleles. For all other contributors, there appears to be an optimal percent of desaturation with corresponding lowest P(RMNE) value. Given this, there are two strategies for implementing mixture analysis with desaturation. First, an overall optimal amount of desaturation can be selected that optimizes the most detections across saturated mixtures – this would be close to 35% to 40% desaturation for these 11 mixtures. Inherent to the problem of saturated mixtures is the chance to match random non-contributing individuals (false positive results). One source of potential false positives is first degree relatives.

The estimated contribution percentages for the eleven mixtures are shown in Figure 4. The lowest contribution percentage of a detected individual was 9%, and the highest contribution percentage of an undetected individual was 7%.

**Figure 4:**
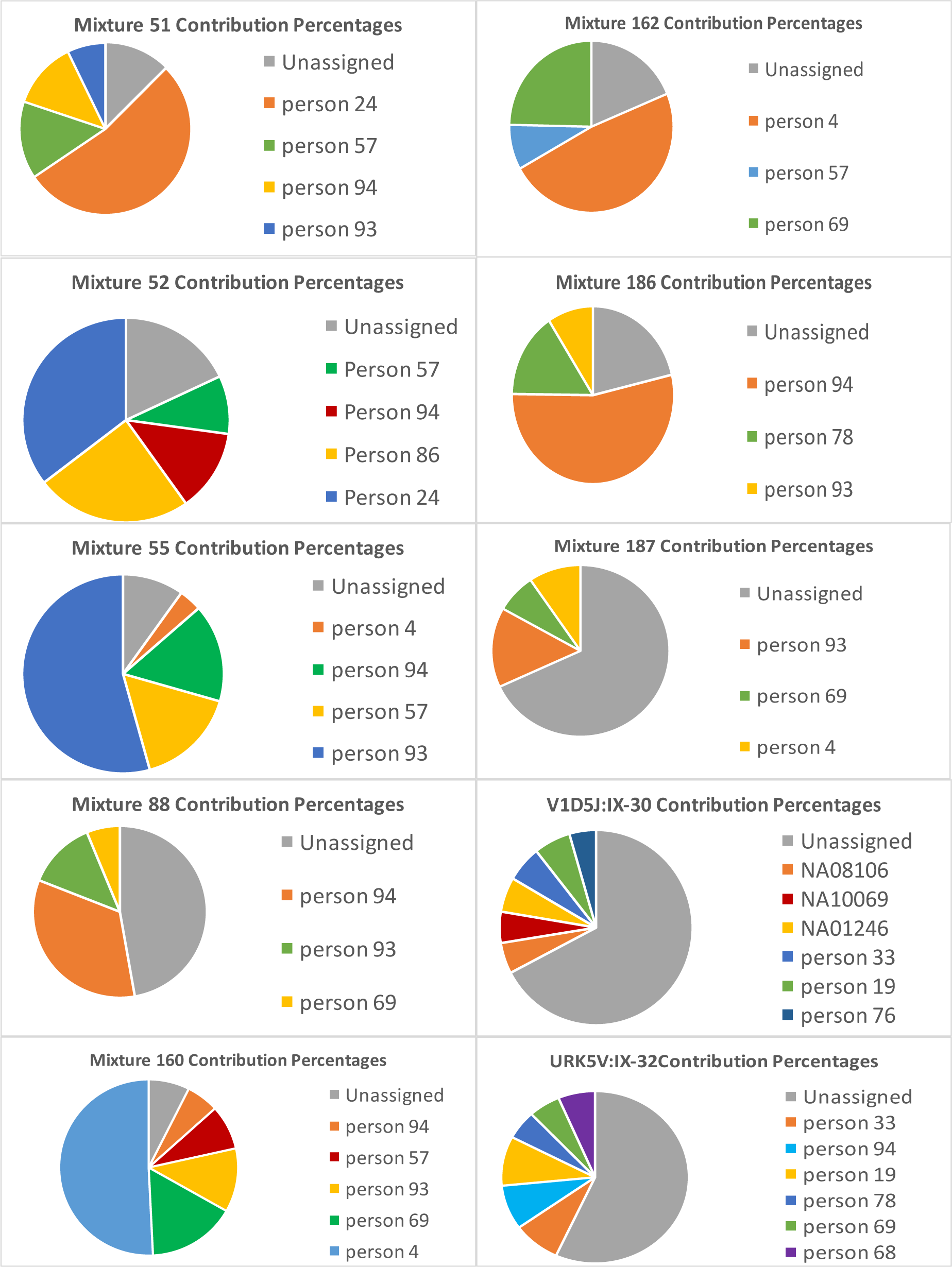
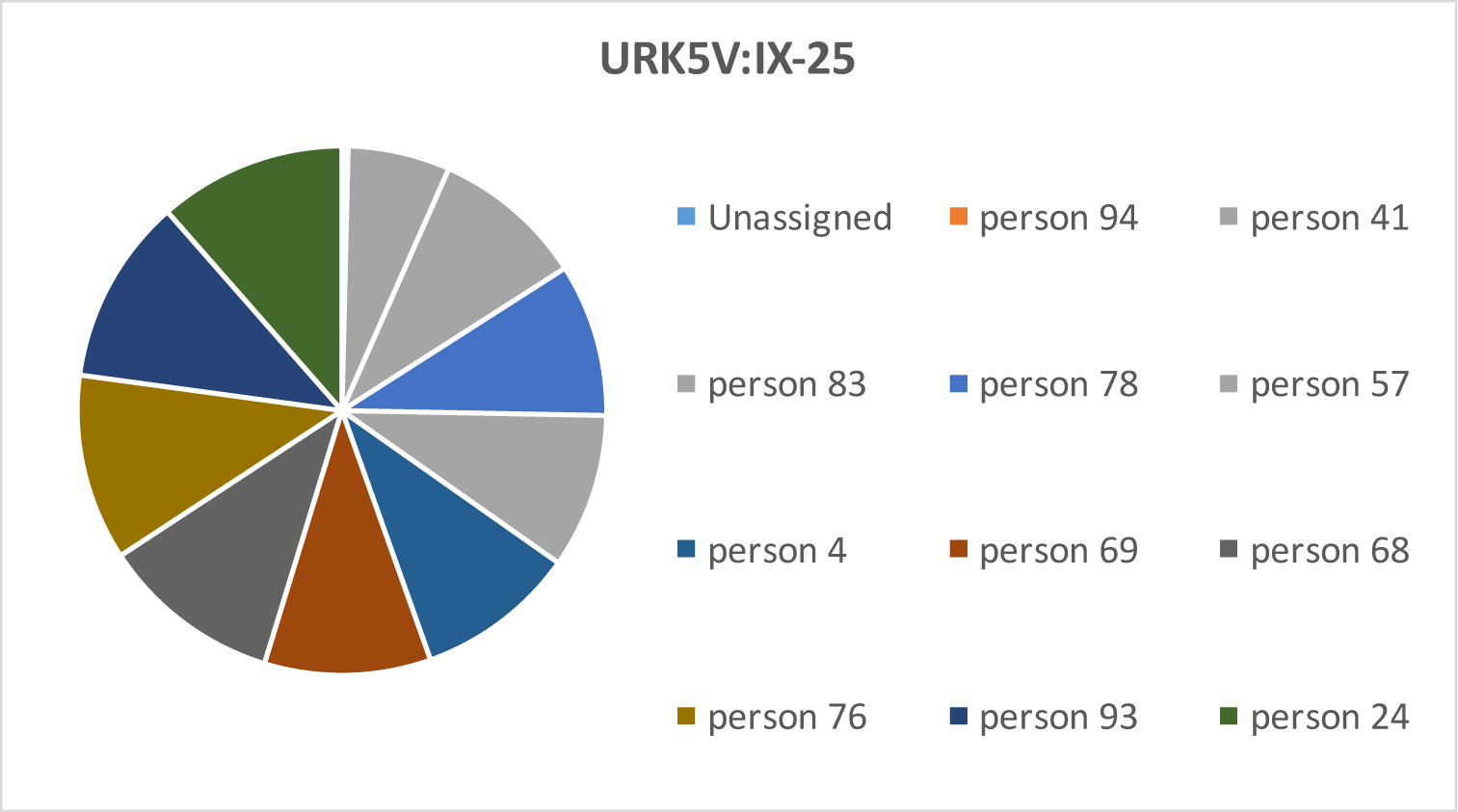
The calculated contribution percentages of individuals identified in the 11 mixtures as well as the portion of the minor alleles that could not be attributed to any of the detected individuals.

### Defined Equimolar Mixtures

Three defined equimolar mixtures were created with DNA from 10, 15, and 20 individuals to test the upper limits of detection on the panel of 2,655 SNPs. Mixture analysis by FastID does detect reference matches, but all matches are filtered by quality filters for saturated mixtures. Mixture analysis by FastID on the TranslucentID desaturated sub-mixtures identifies 9 of 10 (90%), 8 of 15 (53%), and 10 of 20 (50%) of the DNA contributors. This illustrates the value of the combination of mixture desaturation with mixture analysis on saturated mixtures. It would be valuable to test this approach with a larger number of SNP loci in the future; linkage disequilibrium between loci will be an important consideration for the design and optimization of larger SNP panels.

## Conclusions

DNA mixtures with multiple contributors are a challenge for DNA forensics. A major challenge of mixture analysis is the saturation of the panel with DNA from multiple contributors. We introduce the combination of mixture desaturation with TranslucentID and mixture analysis to illustrate the utility of mixture analysis of forensic samples with mixture SNP panels.

## Acknowledgements

DISTRIBUTION STATEMENT A. Approved for public release. Distribution is unlimited.

This material is based upon work supported under Air Force Contract No. FA8702-15-D-0001. Any opinions, findings, conclusions or recommendations expressed in this material are those of the author(s) and do not necessarily reflect the views of the U.S. Air Force.

